# Deciphering *Brassica* plant defence responses to cabbage white butterfly egg-associated molecular patterns

**DOI:** 10.1101/2021.03.29.437462

**Authors:** Lotte Caarls, Niccoló Bassetti, Femke van Doesburg, Patrick Verbaarschot, Joop J.A. van Loon, M. Eric Schranz, Nina E. Fatouros

**Affiliations:** Biosystematics Group, Wageningen University & Research, Wageningen, The Netherlands; Laboratory of Entomology, Wageningen University & Research, Wageningen, The Netherlands

**Keywords:** Plant-insect interactions, hypersensitive response, egg-associated molecular patterns, oviposition-induced plant responses, *Pieris*, *Brassica*

## Abstract

*Brassica* plants activate a strong hypersensitive response (HR)-like necrosis underneath eggs of cabbage white butterflies, but their molecular response to eggs is poorly understood. Here, we developed a method to generate egg wash to identify potential insect egg-associated molecular patterns (EAMPs) inducing HR-like necrosis. We found that egg wash, containing compounds from *Pieris* eggs, induced a similar response as eggs. We show that wash of hatched eggs, of egg glue, and of accessory reproductive glands (ARG) that produce this glue, also induced HR-like necrosis, whereas removal of the glue from eggs resulted in a reduced response. Eggs of *Pieris* butterflies induced callose deposition, production of reactive oxygen species and cell death in *B. nigra* and *B. rapa* leaf tissue, also in plants that did not express HR-like necrosis. Finally, only washes from *Pieris* eggs induced defence genes and ethylene production, whereas egg wash of a generalist moth did not. Our results indicate that EAMPs are in the egg glue and that the response in *B. nigra* is specific to *Pieris* species. Our study expands knowledge on the *Brassica*-*Pieris*-egg interaction, and paves the way for identification of EAMPs in *Pieris* egg glue and corresponding receptor in *Brassica* spp.

## Introduction

Plants rely on an immune system that regulates the perception of attackers and subsequent activation of inducible defences (Wilkinson et al., 2019). Perception involves detection of pathogen-derived effector proteins and of molecular patterns, which can derive from different organisms, such as microbes (MAMPs) or herbivores (HAMPs) (Gust, Pruitt, & Nürnberger, 2017; Stahl, Hilfiker, & Reymond, 2018; Van der Burgh & Joosten, 2019). Perception is followed by an early signaling cascade including rapid ion-flux changes and production of reactive oxygen species (ROS) (Couto & Zipfel, 2016). Induced defences include reinforcement of extracellular barriers, for example by callose deposition, the production of antimicrobial or insecticidal metabolites and proteins, and a localized, rapid cell death response, the hypersensitive response (Balint-Kurti, 2019; Campos, De Souza, De Oliveira, Dias, & Franco, 2018; Couto & Zipfel, 2016; Cui, Tsuda, & Parker, 2015). Up to now, most in-depth studies on activation of the plant immune system have been performed on interactions with plant pathogens, while pattern recognition receptors for HAMPs are just starting to be discovered (Erb & Reymond, 2019; Steinbrenner et al., 2020).

The detection by plants of herbivore eggs deposited on plant tissue, is remarkable, as eggs are immobile and seemingly harmless structures. However, insect eggs turn into feeding larvae and thus pose a threat to the plant (Hilker & Fatouros, 2015, 2016). Upon detection, plants can mount defences against eggs that range from plant-mediated desiccation of eggs, egg dropping, egg crushing and the production of ovicidal toxins. Eggs of cabbage white butterflies (*Pieris* spp) trigger necrotic lesions in leaves of the black mustard, *Brassica nigra* that can result in egg-killing by desiccating and/or dropping off singly laid *Pieris* eggs (Griese, Dicke, Hilker, & Fatouros, 2017; Shapiro & DeVay, 1987). As the phenotype resembles a HR it was termed hypersensitive response-like (“HR-like”) (Fatouros et al., 2012). Oviposition by *Pieris* butterflies has been shown to induce HR-like necrosis in several other plants of the Brassicaceae family, although the severity of the response varies between, and within species (Griese et al., 2021; Groux, Gouhier-Darimont, Kerdaffrec, & Reymond, 2020; Pashalidou, Fatouros, Van Loon, Dicke, & Gols, 2015).

Besides a HR-like necrosis, eggs of different insect species induce immune responses similar to pattern-triggered immunity (PTI), including callose deposition, accumulation of ROS and SA, and transcriptome changes of several defence genes, including *PR1* (Bruessow, Gouhier-Darimont, Buchala, Metraux, & Reymond, 2010; Gouhier-Darimont, Schmiesing, Bonnet, Lassueur, & Reymond, 2013; Little, Gouhier-Darimont, Bruessow, & Reymond, 2007; Lortzing, Kunze, Steppuhn, Hilker, & Lortzing, 2020; Reymond, 2013). Expression of *PR1* was also induced in leaves of *B. nigra* underneath *P. brassicae* and *P. rapae* eggs (Fatouros et al., 2015; Fatouros et al., 2014). Whether other brassicaceous species that are natural hosts of *Pieris* spp., including *B. nigra* and *B. rapa*, respond with a general immune response, including ROS, callose and expression of different defence genes, to insect eggs, and whether there is genetic variation for this response, is unknown.

Species of the Brassicaceae family have co-evolved with their specialists pierid butterflies for millions of years in an arms-race (Edger et al., 2015; Wheat et al., 2007). In a recent study, we suggest that this arms-race has led to the evolution of the HR-like necrosis in *Brassica* plants as an egg-killing trait specifically induced by eggs from brassicaceous-feeding *Pieris* and *Anthocharis* species (Griese et al., 2021). Neither eggs from non-brassicaceous-feeding butterflies, nor from cabbage moths, *Mamestra brassicae* and *Plutella xylostella* induced a necrosis (Griese et al., 2021). On the other hand, eggs of different species were shown to induce a general immune response in *A. thaliana*. Egg extract made from generalist herbivore *Spodoptera littoralis* and from *Drosophila melanogaster* induced *PR1* expression (Bruessow et al., 2010). It is not clear, whether defence responses to eggs observed in Brassicaceous species are general responses that are activated to all insect eggs, or whether they result from detection of a specific *Pieris* EAMP.

So far, few studies have identified EAMPs that activate defence against insect eggs (Hilker & Fatouros, 2015; Reymond, 2013; Stahl et al., 2018). In some studies, secretions surrounding eggs were sufficient to elicit defence responses in plants and a few elicitors have been isolated from these secretions (Hilker, Stein, Schröder, Varama, & Mumm, 2005; Salerno, De Santis, Iacovone, Bin, & Conti, 2013; Tamiru et al., 2011). In *P. brassicae*, egg-enveloping secretions are produced by the accessory reproductive gland (ARG) and form a glue-like structure between the eggs and leaves (Beament & Lal, 1957; Fatouros et al., 2012). Treatment of *Brassica* plants with the ARG has been shown to induce HR-like necrosis and plant chemical cues attracting egg parasitoids, and to prime plants for future larval attack (Fatouros et al., 2008; Fatouros et al., 2015; Fatouros et al., 2009; Paniagua Voirol et al., 2020). Anti-aphrodisiacs transferred from the male during mating were shown to be present in minute amounts in the ARG secretion, and were suggested as potential elicitors (Fatouros et al., 2008; Fatouros et al., 2009). However, glands from unmated females induced HR-like, and therefore another female-derived elicitor is likely to play a role (Fatouros et al., 2015). In *A. thaliana*, egg-derived phosphatidylcholines were recently found to induce H_2_O_2_, SA and trypan blue staining (Stahl et al., 2020). It is thus still unclear which compounds from *Pieris* eggs are detected by *Brassica* spp. plants and result in HR-like activation, and whether these reside in the eggs themselves or in the secretions surrounding the eggs.

In this study, we developed and implemented a new method to screen compounds from *Pieris* spp. eggs and egg-enveloping secretions. We specifically addressed: 1) where the EAMP of HR-like necrosis originates from, 2) the cellular and molecular response of two *Brassica* plants and 3) the specificity of these responses by comparing eggs and egg washes of *Pieris* with the generalist moth *M. brassicae*.

## Material and methods

### Plant material and rearing of butterflies

Black mustard (*B. nigra* L.) plants used, originated from plants that were collected near the river Rhine in Wageningen, The Netherlands (N51.96, E05.68). *Brassica rapa* genotypes L58, R-o-18, and RC-144 were obtained from the Laboratory of Plant Breeding (WUR). Plants were grown in a greenhouse (18 ± 5 °C, 50–70% RH, L16: D8) and were used when three to five weeks old.

*Pieris brassicae* L. (Lepidoptera: Pieridae) was reared on Brussels sprouts plants (*Brassica oleracea* var. *gemmifera* cv. Cyrus) in a climate room (21 ± 1 °C, 50–70 % RH, L16: D8). Virgin adult females were obtained by isolating female butterflies immediately after eclosion. Otherwise, twenty females and males could mate in a large cage (60 x 60 x 90 cm) and females were used for oviposition in experiments or oviposition on paper for egg wash production. The cabbage moth *Mamestra brassicae* L. (Lepidoptera: Noctuidae) was reared on Brussels sprouts plants in a climate room (21 ± 1 °C, 50–70% RH, L16: D8).

### Preparation of egg washes

*Pieris brassicae* eggs needed for preparing the egg wash were collected on filter paper pinned underneath a *B. oleracea* leaf in a large cage containing twenty mated females (for details see results section). To collect unfertilized eggs for egg wash, a similar setup was used, except paper was pinned underneath a *B. nigra* leaf of a plant left in a cage with ten virgin butterflies.

To obtain egg-enveloping secretions for wash of egg glue, *P. brassicae* eggs were collected as above, counted and then carefully removed from the paper using a brush. The spots of secretions underneath eggs were then cut out and washed overnight in 1 mL 20 mM MES buffer (pH 5.7) per the equivalent of 400 eggs. A wash of the pieces of the same filter paper without secretions was used as control. To remove egg-enveloping secretions from eggs, eggs on paper were submersed in either a solution of 1 % bleach and 2 % Tween-20 or in 250 mM NaPO_4_ pH 9.0 for 30 minutes. After treatment, eggs were rinsed with MQ and MES buffer and then washed overnight in 20 mM MES buffer pH 5.7. Control eggs were submersed in MES for 30 minutes, rinsed in MQ and MES, and then washed. To study the induction by eggs of different ages, eggs were collected as above on paper, and then kept at room temperature until they were washed at the end of each day, until they hatched. *Pieris brassicae* eggs hatched 6-7 days after oviposition. After hatching, young caterpillars were carefully removed using a brush. Empty eggshells and associated secretions (on paper) were washed overnight.

Paper sheets with *M. brassicae* oviposited eggs were obtained from the Laboratory of Entomology (WUR). To obtain egg wash, eggs were counted, cut out, and washed including paper, in 1 mL 20 mM MES buffer (pH 5.7) per 400 eggs. The wash was pipetted off the next morning and frozen until use.

### Dissection of reproductive tract

For dissection of tissues from the reproductive tract (the accessory reproductive gland (ARG), bursa copulatrix or eggs from the ovarian tubes (“ovarian eggs”) (Supplementary Figure S1)), mated *P. brassicae* females were obtained by pairing a virgin female and virgin male one day after eclosion. Two days after mating, females were killed and dissected. ARGs, bursa copulatrix and ovarian eggs were dissected from mated and virgin *P. brassicae* females (both 3-4 days after eclosion) under a stereomicroscope (optical magnification 20×) in 20 mM MES buffer. Dissected structures were washed overnight in 20 mM MES pH 5.7 using 50 µL per each ARG, 100 µL buffer per each BC or 5 µL buffer per egg. Solution was pipetted off the next morning and frozen until use.

### Treatment of plants with egg wash and egg deposition by butterflies

For all experiments testing the effects of treating plants with egg washes, 10 µL egg wash was pipetted on the abaxial side of the fourth or fifth emerged *B. nigra* leaf of 3-4 weeks old plants. Symptoms induced by egg wash were scored four days after treatment. To quantify severity, a scoring system was used from 0-4 (Figure 1E).

**Figure 1.**
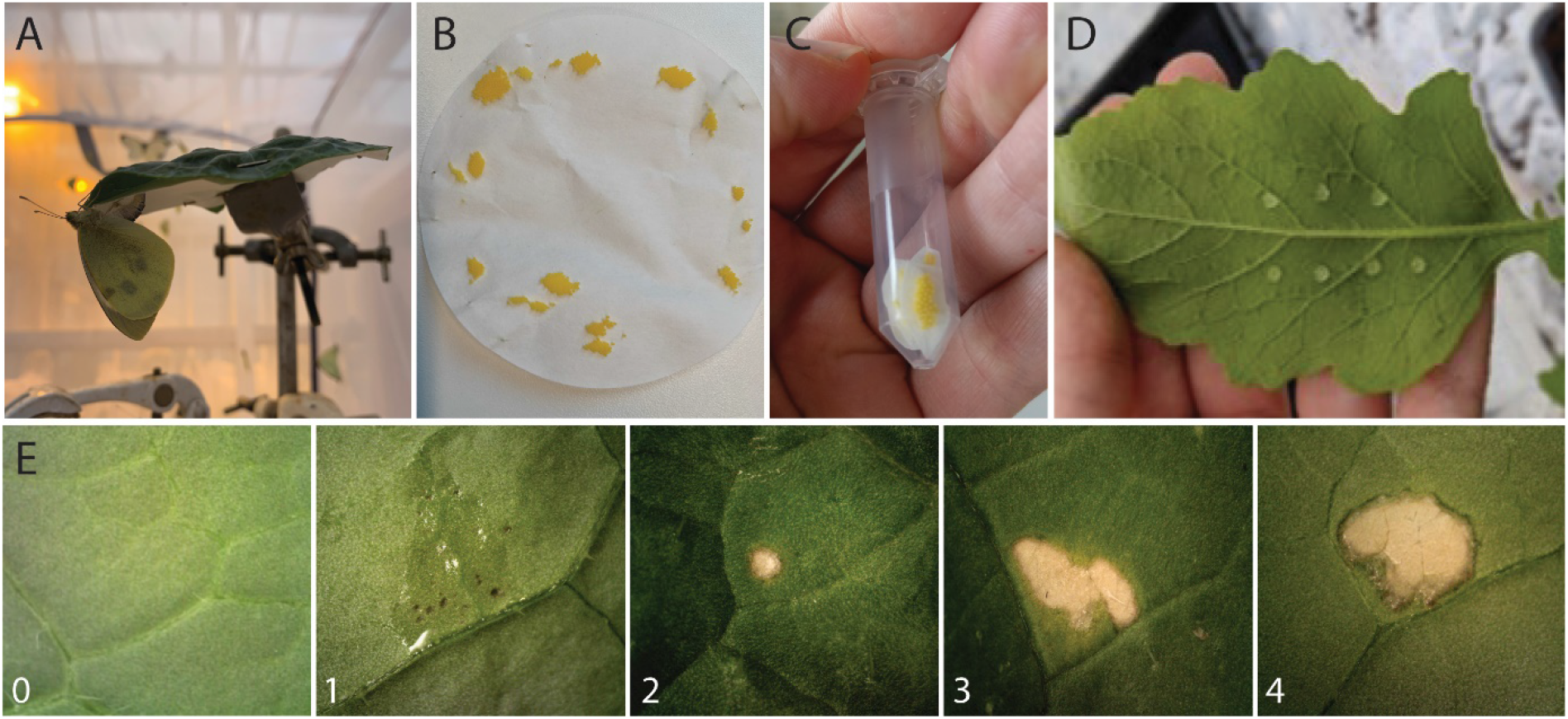
Collection of eggs, preparation of egg wash and scoring of HR-like symptoms. A) Setup in a greenhouse with *P. brassicae* butterfly depositing eggs on paper pinned underneath *B. oleracea* leaf. B) Picture of clutches of *P. brassicae* eggs as deposited on paper. C). Eggs are cut out of paper and washed overnight in an Eppendorf tube in 1 mL buffer/400 eggs. D) Response of leaves is tested by applying drops of 10 µL egg wash on the abaxial side of the leaf, where eggs are usually deposited by *P. brassicae*. E) Representative pictures of classes used to score severity of visual symptoms induced by egg wash: 0: no visual response. 1: brown spots underneath eggs or egg wash spot, only visible at abaxial side leaf. 2: necrosis also visible at adaxial side of leaf, spot smaller than 2 mm diameter, 3: necrosis the size of egg wash spot, and 4: spreading lesion beyond spot of treatment. Score 0 and 1 are classified as “non-HR-like”, score 2-4 are classified as “HR-like”.

For experiments with oviposition on plants, one *P. brassicae* butterfly was placed in a cage with a *B. nigra* or *B. rapa* plant and removed when the required number of eggs were laid. HR-like necrosis was scored three or four days after oviposition, using the same scoring system as with egg wash. For experiments with *M. brassicae*, female moths were placed together with a *B. nigra* plant in a cage to allow egg deposition overnight and removed in the morning.

### Expression of genes by real-time qRT-PCR

For measurement of gene expression, plants were treated with egg wash or oviposited by butterflies as above. For comparison of induction of gene expression by eggs and egg wash, *B. nigra* plants were induced by either 10 µL of egg wash, or an egg clutch of 10 eggs. Samples were then harvested immediately, or after 3, 6, 24 and 48 h by taking six leaf discs (Ø 6 mm) directly next to the eggs or the spot of egg wash treatment. For each timepoint, four plants were sampled individually and considered biological replicates. *B. rapa* plants received three single eggs on a single leaf of each plant, and leaf discs (Ø 6 mm) were harvested next to the eggs, immediately, or after 3, 6, 24 and 96 h. For each timepoint, three biological replicates were used. To compare the induction of genes between *P. brassicae* and *M. brassicae* egg wash, plants were treated with 10 µL of either egg wash or a control MES buffer. Samples were then harvested after 24 hours by taking six leaf discs (Ø 6 mm) directly next to the spot of egg wash treatment. Four plants were used for each treatment as biological replicates. A standard protocol was used for RNA isolation, cDNA synthesis and q-RT-PCR (Supplemental Data).

### Production of ethylene

To measure the plant production of ethylene, leaves of untreated plants were harvested. Later, the same plants were used to assay induction of the HR-like response by *P. brassicae* egg wash. To compare ethylene production between *B. nigra* plants with contrasting HR-like response, for each HR-like response (no or yes), ten plants were used. For *B. rapa*, three plants of each genotype were used. Ethylene production was measured as published (Oome et al., 2014), five hours after incubation of three leaf discs of 3 mm in either 400 µL of 20 mM MES pH 5.7 or 400 µL egg wash or egg wash diluted four times in 20 mM MES. Ethylene was analysed on a Focus gas chromatograph (Thermo Electron S.p.A., Milan, Italy) equipped with an FID detector and a RT-QPLOT column, 15 m × 0.53 mm ID (Restek, Bellefonte, PA, USA). The system was calibrated with a certified gas of 1.01 μL L−1 (1 ppm) ethylene in synthetic air (Linde Gas Benelux B.V., Schiedam, The Netherlands).

### Histochemical staining

For histochemical staining, plants were used for egg deposition by *P. brassicae* and samples were taken 24, 48 and 72 hours after oviposition by taking a 10 mm diameter leaf disc of the area surrounding the eggs or egg wash. Pictures of the leaf discs were taken with a Dino-Lite digital microscope (AnMo Electronics Corporation) before the eggs were carefully removed. Nitroblue tetrazolium (NBT; Sigma) was used to stain superoxide radical O_2_^•−^. For this, leaf discs were submersed in 2 ml 0.2% NBT and 50 mM sodium phosphate buffer (pH 7.5) and samples were incubated 30 to 60 minutes in the dark. For visualization of cell death, leaves were submersed in 0.08 % trypan blue solution (Sigma), overnight. For staining of callose, destained leaf discs were submersed in 0.01 % aniline blue in 150 mM K_2_HPO_4_ and imaged after at least 2 hours of incubation using a DAPI filter on a fluorescence microscope with NIS elements AR 2.30 software.

### Data analysis

All data analysis was carried out in R (R Core Team, 2020). The occurrence of HR-like necrosis was analysed with a generalized linear model (GLM) with a binomial error distribution. The response of plants to each treatment was considered as a binomial response: either non-HR (score 0 or 1) or HR (score 2, 3, and 4) and different treatments were included as categorical fixed factors. When overall differences were found, pairwise differences between factors were tested. HR severity was considered as the score of symptoms induced, and differences in mean HR severity were tested using Kruskal-Wallis test, followed by Wilcoxon Rank Sum test with Benjamini-Hochberg correction. For gene transcription data, ΔΔCq values were used for statistical analysis. To compare eggs and egg wash, data were analysed with one-way ANOVA for the two treatments (eggs or egg wash) independently, followed by Dunnett’s test to compare all timepoints to the 0h timepoint. The differences between treatments were tested for each timepoint with Student’s t-tests. To compare gene transcription data between plants treated with either *Pieris brassicae* or *Mamestra brassicae*, ΔΔCq values were used for statistical analysis with one-way ANOVA, followed by Tukey post-hoc tests. Differences in mean ethylene produced (ppm) after treatments or between plants were tested using the Kruskal-Wallis test, followed by the Wilcoxon Rank Sum test with Benjamini-Hochberg correction.

## Results

### Development of egg wash methodology

We developed a method to isolate *P. brassicae* eggs and egg-enveloping secretions, and dissolve potential EAMPs (Figure 1). *Pieris* butterflies are specialists on brassicaceous plants, and oviposition is stimulated by chemicals present on the leaf surface, i.e. glucosinolates, detected by sensilla present on the butterflies’ tarsi (Städler & Reifenrath, 2009; Van Loon et al., 1992). Our aim was to collect compounds from the surface of eggs and avoid molecules from the leaves. We therefore constructed a setup that included an oviposition stimulus but excluded molecules from the plant in the subsequent collection of eggs.

To this end, a round filter paper was attached to the abaxial side of a detached *B. oleracea* leaf, exactly covering the leaf surface. By perceiving the oviposition stimulus from the leaf surface via the tarsi, oviposition is stimulated: the female curves her abdomen to the abaxial side and in this way, deposits her eggs on the pinned paper (Figure 1A). Oviposition by twenty butterflies in a cage resulted in the collection of several egg clutches, up to a few hundred eggs per paper (Figure 1B). Egg clutches laid on the paper were cut out and submersed in buffer overnight without disturbance (Figure 1C). In this way, eggs remained undamaged, viable, and neonate larvae could still hatch from them (L. Caarls, unpublished data). The solution (egg wash) was pipetted into a new tube the next morning.

When used for treatment of plants, egg wash is pipetted, like eggs are deposited, on the abaxial side of a leaf (Figure 1D). The necrosis induced by egg wash becomes visible 1-2 days after treatment, and is scored four days after treatment, in five classes (Figure 1E).

### Egg wash induced a plant response similar to the response to eggs

When compared visually, the treatment of leaves with egg wash induced a similar HR-like necrosis as deposition of eggs by butterflies (Figure 2A). Symptoms induced by eggs or egg wash were scored after four days, and HR-like frequency (proportion of plants showing HR-like) and HR-like severity (mean score of induced symptoms) were compared between the two treatments. HR-like frequency after oviposition by eggs or treatment with egg wash did not differ. On average, egg wash induced slightly higher HR-like severity than eggs (Figure 2B, Supplementary Table S2).

**Figure 2.**
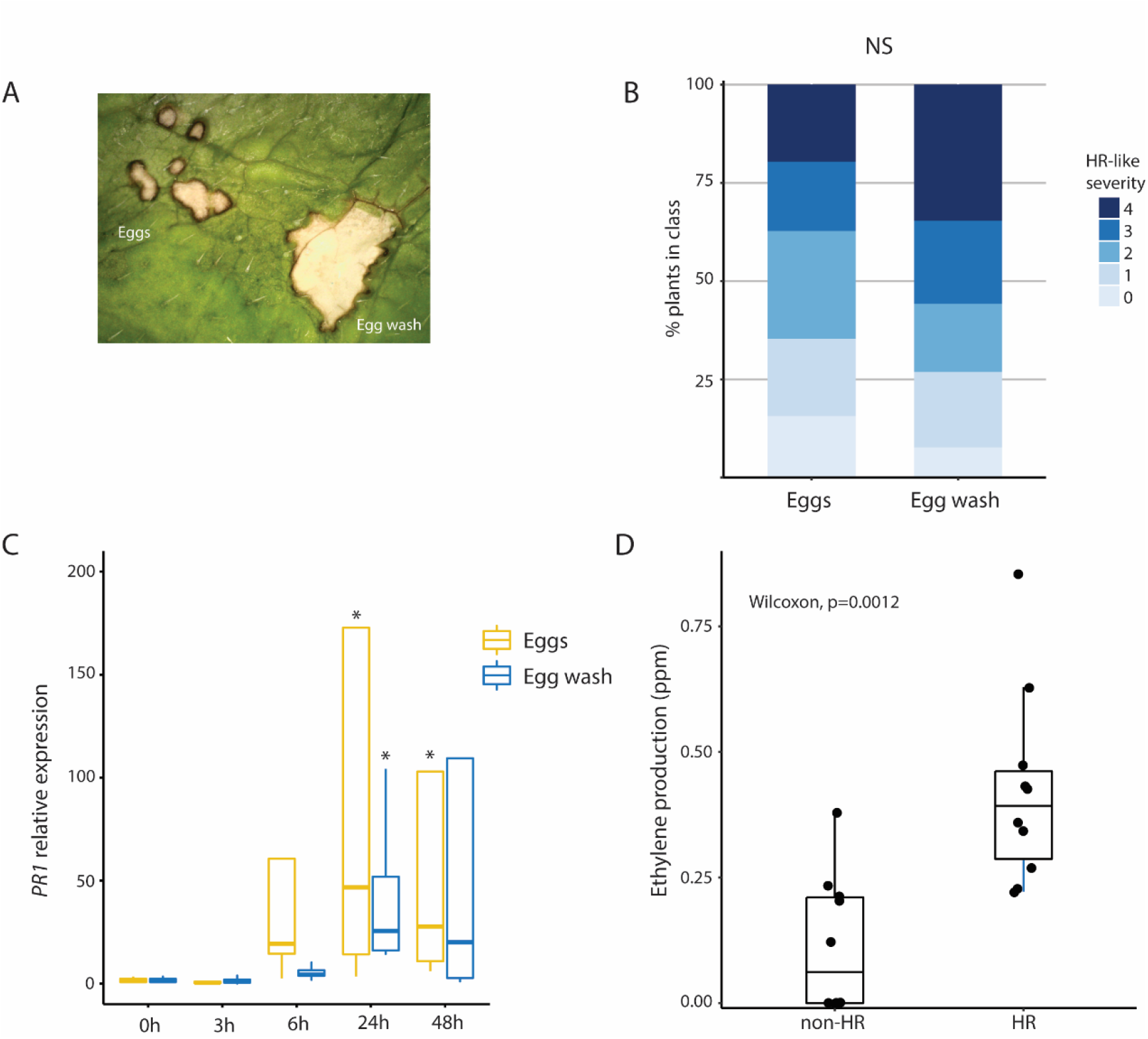
Plant responses induced by *Pieris brassicae* eggs and egg wash in *B. nigra*. A) Picture of responses induced by eggs and egg wash next to each other on leaf (microscopic image): both eggs and egg wash induce HR-like necrotic spots. B). Quantification of severity of symptoms induced by egg wash and eggs. For classes score, see Figure 1E. NS = no significant difference (Kruskal-Wallis: H = 3.44, df = 1, P = 0.06). C). *PR1* expression in plants after egg deposition (yellow box plots) or treatment with egg wash (blue box plots). Asterisks indicate significantly higher expression at the timepoint compared to the 0 h timepoint (ANOVA followed by Dunnett’s test, P < 0.05). D) Ethylene production in parts per million (ppm) by plants treated with egg wash. Ethylene production by plants that show HR to egg wash was significantly higher compared to plants that did not (Wilcoxon rank sum test, P = 0.0012).

We then measured expression of *PR1* in leaves oviposited on by *P. brassicae* butterflies or treated with egg wash. *PR1* was significantly upregulated both after oviposition and after treatment with egg wash. *PR1* expression increased after 6 hours and was significantly induced in plants 24 hours after treatment with egg wash, and 24 and 48 hours after egg deposition, compared to expression level prior to treatment (Supplementary Table S3). No significant differences in *PR1* expression were found between plants treated with eggs or egg wash (Figure 2C, Supplementary Table S3).

Next, we tested whether egg wash induces ethylene. *Brassica nigra* leaves responded with ethylene production after incubation with egg wash for 5 hours compared to incubation with control MES buffer (Supplementary Table S4). There was a significant difference in ethylene produced between plants with contrasting responses. Plants responding with stronger HR-like necrosis, produced a significantly higher amount of ethylene after incubation with egg wash than plants with no HR-like necrosis (Figure 2D, Supplementary Table S4). Similarly, in the *B. rapa* responsive accession L58, incubation with *P. brassicae* egg wash also resulted in ethylene production (Supplementary Figure S2A). These results suggest that there is an early detection response in plants after contact with egg wash that will ultimately lead to cell death in some plants.

### EAMP derived from female accessory reproductive glands and not from inside the egg

To study if eliciting molecules originate from inside the eggs or from egg-enveloping secretions, we studied if eggs and secretions lose their HR-like eliciting activity with aging, or when eggs hatch. There was no significant difference in eliciting activity of eggs of increasing age, from one-day-old (the egg age at which egg wash usually is made), to five-day-old eggs, although HR-like severity decreased slightly. When caterpillars hatched, and only (6-day-old) eggshells and secretions remained on paper and were washed, the wash still induced HR-like symptoms (Figure 3A Supplementary Table S5).

**Figure 3.**
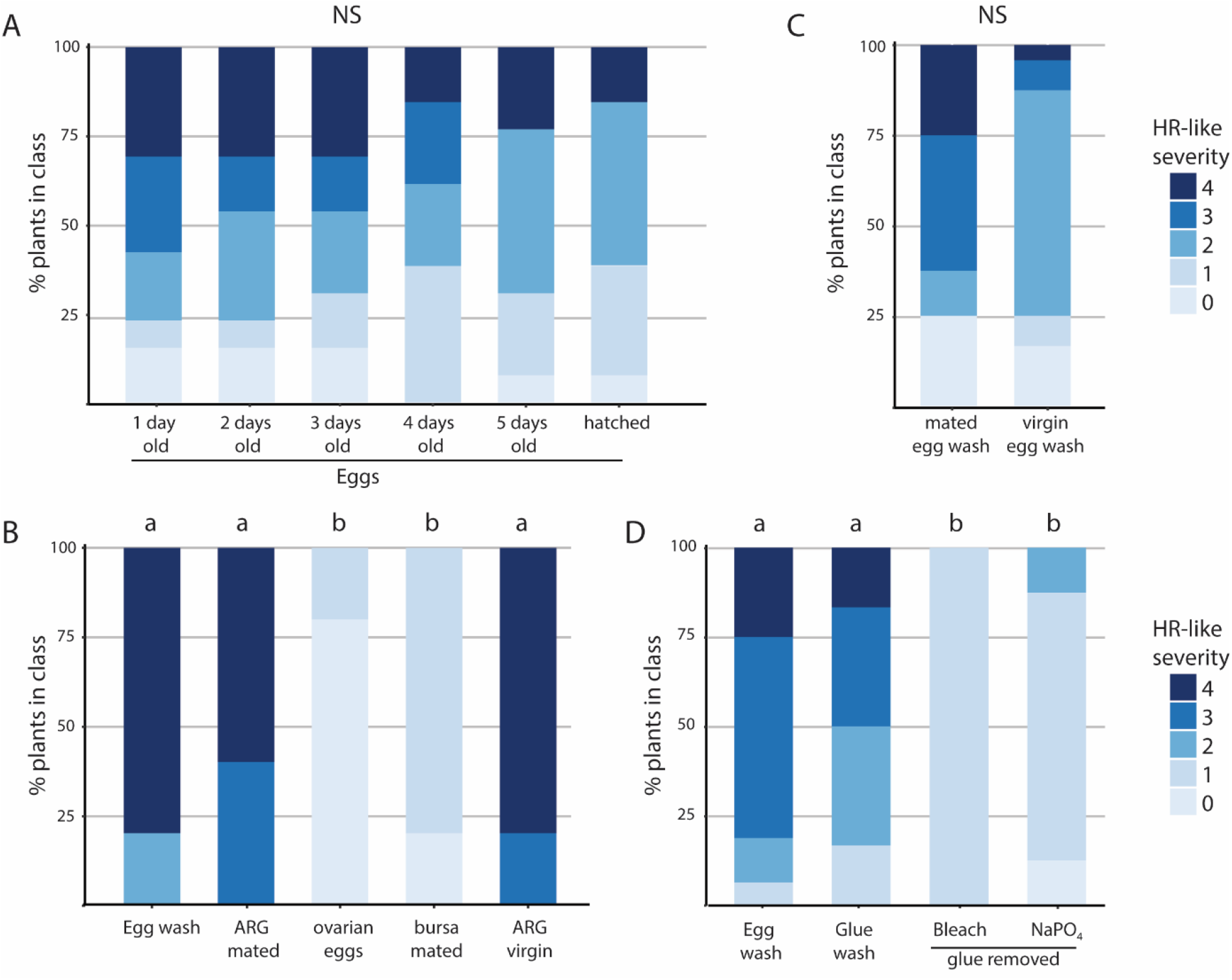
EAMP found in egg wash is in the glue, produced in the ARG and is female-derived. A-D). Severity of HR-like symptoms induced in plants by the different washes. For classes score, see Figure 1E. A) Egg washes made of eggs of different ages or eggshells and egg glue remaining on filter paper. B) Egg wash of dissected structures of reproductive tract. Different letters indicate significant differences in HR severity (Kruskal-Wallis, H = 24.06, df = 2, P < 0.001). C) Egg wash of eggs of mated females versus virgin females. D) Wash of eggs, glue alone (in filter paper) or wash of eggs with egg-enveloping secretions removed. For glue removal, two treatments were used, either a 1% bleach wash or wash with NaPO_4_. Different letters indicate significant differences in HR severity (Kruskal-Wallis: H = 26.60, df = 3, P < 0.001).

In butterfly females, eggs are produced in the ovaries, pass through the common oviduct and are fertilized by sperm released from the bursa copulatrix (BC) into the vagina. Before being expelled through the ovipore, the eggs are covered by secretions released from the accessory reproductive gland (ARG), a paired gland that contains egg-enveloping secretions and cement to glue eggs to leaves (Supplementary Figure S1). We tested wash made from dissected structures of the female reproductive tract of *P. brassicae*. A wash of dissected ARGs induced necrotic spots, similar to the positive control *P. brassicae* egg wash (Figure 3B). On the contrary, neither a wash of unfertilized but mature eggs dissected from the ovary (‘ovarian eggs’) nor a wash of the BC, induced symptoms (Figure 3B). HR-like severity was significantly higher in plants treated with egg wash or a wash of ARG, compared to wash of ovarian eggs or bursa copulatrix (Supplementary Table S5).

It was then investigated, if the inducing molecules in the ARG secretions are of female or male origin. A wash from unfertilized, deposited eggs (containing secretions from the ARG) from virgin butterflies induced a similar response as wash from fertilized, deposited eggs of mated females (Figure 3C). There was no significant effect of the mating status of the female (mated or virgin) on the frequency of HR-like necrosis elicited or on HR severity. In addition, the wash of ARGs from virgin females induced strong symptoms similar to those of mated females (Figure 3B). These results show that egg fertilization is not necessary for the induction of the HR-like necrosis in *B. nigra*, and that an EAMP resides in the ARG, and is female-derived.

Finally, we tested a wash made from the remaining glue left on filter paper after *P. brassicae* eggs are carefully removed. In addition, the glue enveloping oviposited eggs was removed and eggs without glue were washed. Both egg wash and wash of glue alone induced a severe HR-like necrosis, and the severity was significantly lower when *B. nigra* was treated with a wash of eggs from which the glue was removed (Figure 3D; Supplementary Table S5).

### *Pieris* eggs triggered cellular responses in two *Brassica* species

Next, we investigated cellular responses against *Pieris* oviposition by comparing two plant species, *B. nigra* and *B. rapa*, that differ in the severity of the HR-like response. In *B. rapa*, the visual response to *Pieris* eggs consisted of blackening underneath eggs and some necrosis. In addition, in *B. rapa* the expression of *PR1* in leaves was increased 24 hours and significantly higher 96 hours after egg deposition, compared to plants before treatment (Supplementary Figure S2B).

In *B. nigra*, trypan blue staining revealed cell death in the leaf underneath oviposited eggs (Figure 4A-B). In leaves that show no visual necrosis at the place of oviposition (Figure 4C), trypan blue also clearly stained underneath the eggs (Figure 4D). For *B. rapa*, trypan blue stained cell dead in leaf tissue underneath eggs as well, regardless of the occurrence of a necrosis visible by eye (Supplementary Figure S2C-2H).

**Figure 4.**
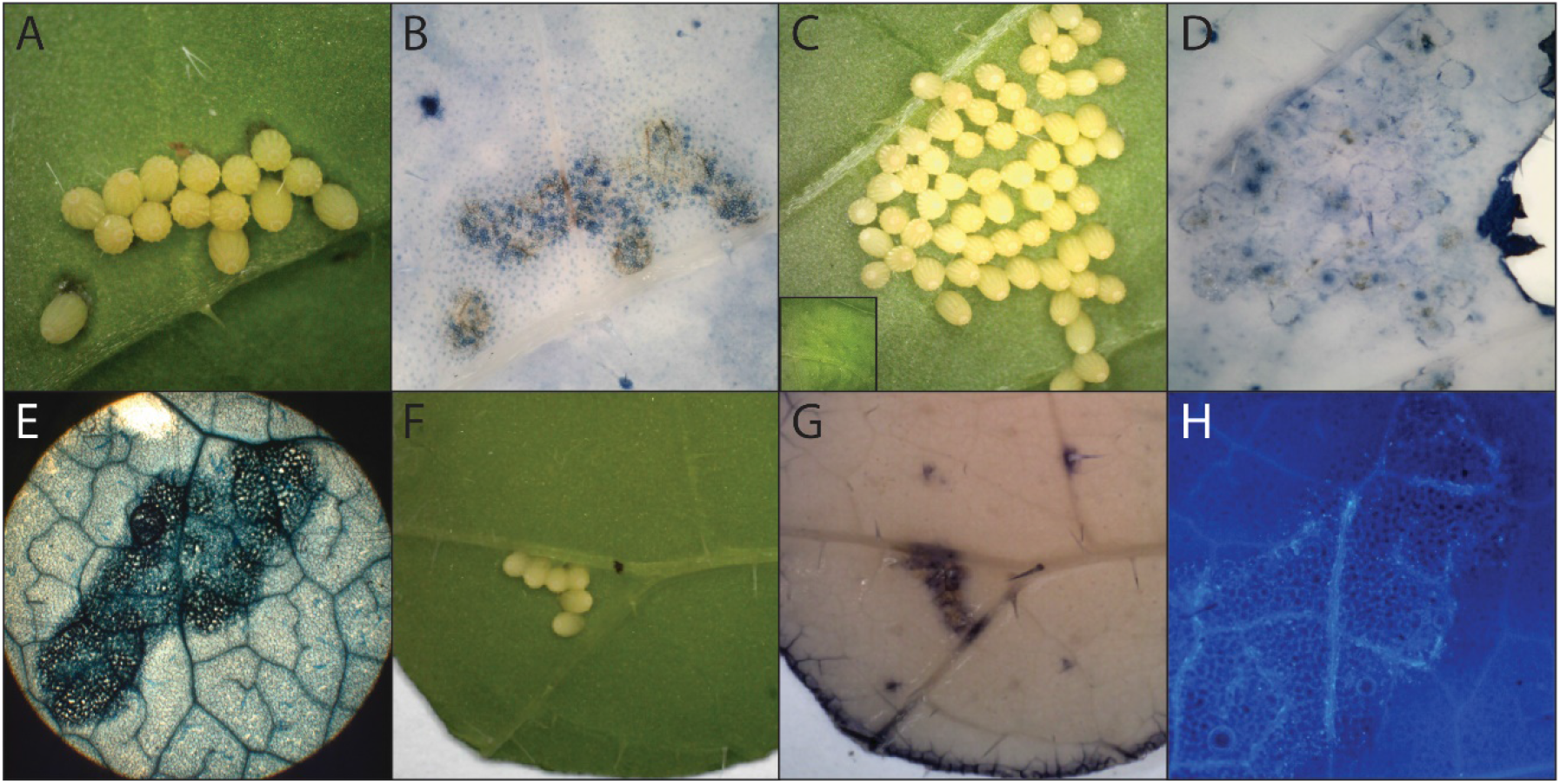
Eggs of *P. brassicae* induce cellular responses in *B. nigra*. A) *B. nigra* leaf 72 h after oviposition. B) Trypan blue staining of leaf shown in (A) showing cell death underneath eggs. C) *B. nigra* leaf with no visible HR-like necrosis underneath eggs 72 h after oviposition. Insert: adaxial side of same leaf showing no visible response. D) Trypan blue staining of leaf shown in C. revealing dead cells underneath eggs. E) Microscopic image of trypan blue stained leaf visualizing egg-clutch shaped stain. F) *B. nigra* leaf 24 hours after oviposition. G) NBT staining of leaf shown in (F) revealing O_2_^•−^ deposition underneath eggs. H) Microscopic image of leaf stained with aniline blue showing callose deposition underneath eggs 24 h after oviposition.

Microscopic investigation of the stains revealed that the stain was shaped as an egg clutch, suggesting the cell death is often isolated to the leaf tissue directly underneath the eggs (Figure 4E). The accumulation of O_2_^•−^ was detected 24 hours after oviposition underneath eggs, both in *B. nigra* and *B. rapa* (Figure 4F and 4G, Supplementary Figure S2I-J). By staining *B. nigra* leaves with aniline blue, callose deposition was found underneath eggs deposited on *B. nigra* (Figure 1H), and surrounding the necrotic spot caused by eggs.

### Moth eggs and egg wash do not induce similar responses as *P. brassicae* in *B. nigra*

To understand whether the cellular and molecular response in *B. nigra* is specific to cabbage white butterfly eggs, we compared responses in *B. nigra* to *P. brassicae* eggs with those to *M. brassicae* eggs and egg wash. Staining of plants showed that plant cells did not die underneath *M. brassicae* eggs as leaves did not stain with trypan blue (Figure 5A and B, compare Figure 4). Further, *M. brassicae* eggs induced O_2_^•−^ production in some plants, but weaker than *Pieris* eggs (Figure 5C and D). While *P. brassicae* egg wash induced the expression of *PR1* after 24 hours, *PR1* expression induced by *M. brassicae* egg wash was not different from the control treatment (Figure 5E, Supplementary Table S5). In addition, incubation with *M. brassicae* egg wash induced low ethylene production, that was significantly lower than that produced after treatment with *P. brassicae* egg wash (Figure 5F).

**Figure 5.**
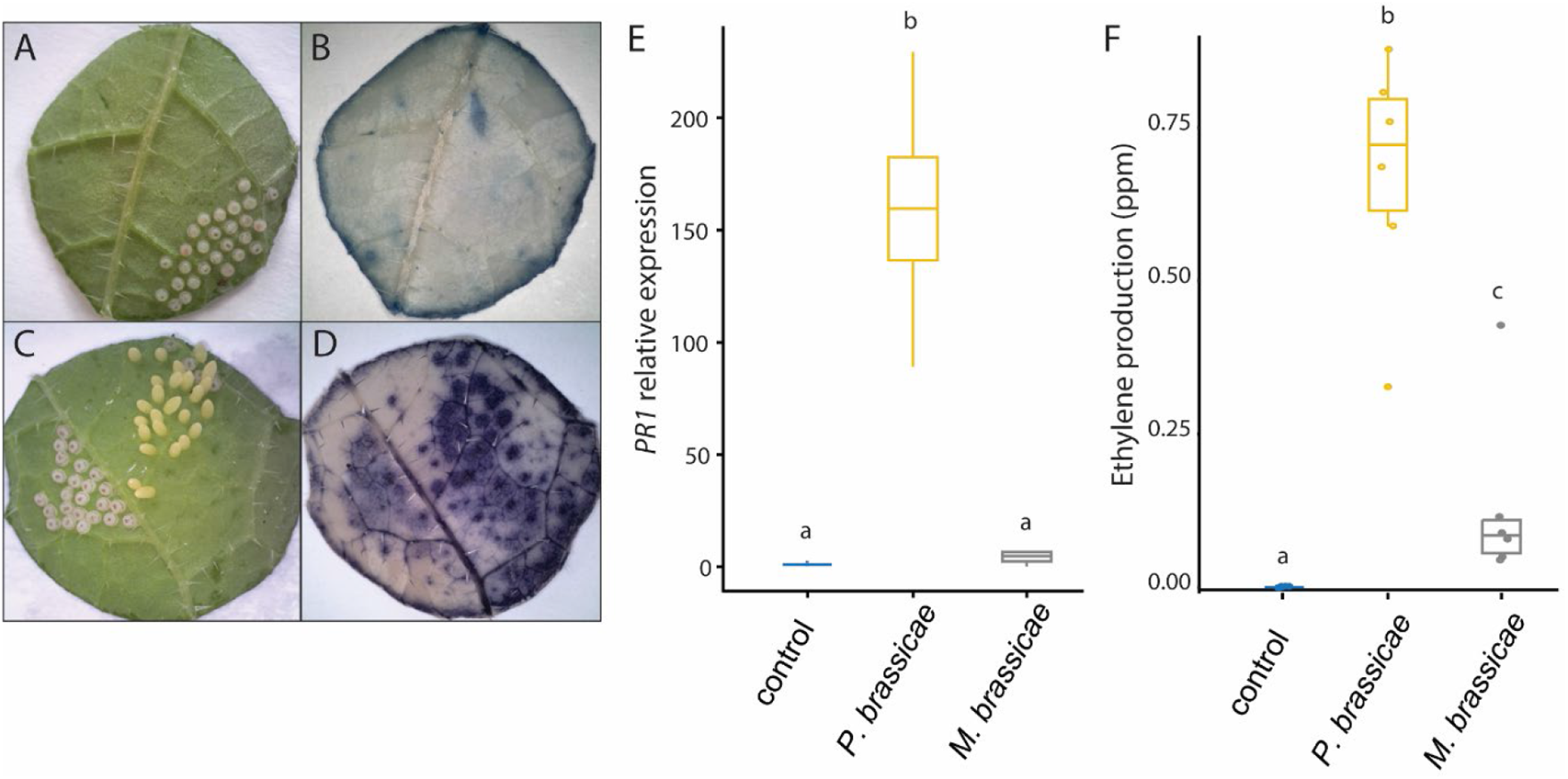
Responses to eggs and egg wash of *M. brassicae*. A) Leaf of *B. nigra* oviposited on by *M. brassicae* moth. B) Leaf stained by trypan blue showing no cell death underneath *M. brassica*e eggs. C) Leaf oviposited with *P. brassicae* (yellow) and *M. brassicae* (white) eggs next to each other. D) O_2_^•−^ production in leaf in C stained with NBT to reveal strong O_2_^•−^ production underneath *P. brassicae* eggs and light underneath *M. brassicae* eggs. E) *PR1* expression in leaf tissue treated with control solution, *P. brassicae* wash or *M. brassicae* wash. Different letters indicate significant differences in mean *PR1* expression, ANOVA followed by Tukey, P < 0.001. F) Ethylene production in *B. nigra* leaf in response to egg washes. Different letters indicate significant differences in mean production of ethylene, pairwise Wilcoxon test, P < 0.01.

## Discussion

In this study, we show that HR-inducing EAMPs are in the egg-enveloping secretions and produced in female ARGs. When plants are treated with egg wash, HR-like necrosis is induced in responsive plants, together with *PR1* expression and ethylene production. In addition, deposition of eggs leads to ROS and callose deposition, also in plants that do not show HR-like necrosis. We find that these phenotypes are shared between two *Brassica* species that vary in the severity of the HR-like response and are specific to eggs and egg wash of the specialist *P. brassicae*.

The increasing knowledge on plant defence mechanisms to eggs of herbivores, suggests that plants can specifically recognize and respond to egg deposition, presumably via the detection of EAMPs. However, very few EAMPs have been identified (Hilker & Fatouros, 2015; Reymond, 2013; Stahl et al., 2018). We have developed a method to obtain and study EAMPs by washing eggs of specialist butterflies. By manipulating the butterflies to lay their eggs on filter paper, compounds from eggs could be isolated without contamination from leaves. The use of egg wash instead of eggs has several other advantages: egg wash can be easily treated to study characteristics of the EAMP, and can be used as a reproducible egg-mimicking treatment in germplasm screenings (Griese et al., 2021). Finally, the use of egg wash allows measurement of other plant defence-related phenotypes that can be used as markers to guide the identification of EAMPs.

When washing eggs, mainly compounds from outside of the eggs and from egg-enveloping secretions are dissolved. We present evidence that at least one *Pieris*-specific EAMP is in these secretions that envelop the eggs: i) wash of glue on filter paper is sufficient to induce the HR-like necrosis, ii) when the secretions are removed from eggs, and eggs are then washed, HR-like necrosis in plants is diminished, and iii) a wash of ARGs, the organs where secretions are produced, is also sufficient to induce HR-like necrosis. As the egg surface and egg-exterior associated secretions are in direct contact with the leaves, it could be expected that plants evolve to detect elicitors in egg-enveloping secretions. We believe it resembles the natural situation of leaf-egg interaction, more so than, for example, crushing of eggs (Bruessow et al., 2010; Little et al., 2007) or crushing of adults (Doss et al., 2000; Y. Yang et al., 2014). Previously, a male-derived anti-aphrodisiac compound transferred during mating to the female ARG, benzyl cyanide, was presented as potential elicitor (Fatouros et al., 2008). We find no evidence for a male-derived elicitor: eggs and ARGs of virgin *P. brassicae* butterflies induced HR-like necrosis in an equal manner as mated butterflies. We thus hypothesize that at least one HR-inducing EAMP is present in the egg-enveloping secretions and ARGs, and is female-derived. Chemical analysis of egg wash and glands will be carried out to identify this EAMP.

Recently, phosphatidylcholines (PCs) were identified as EAMPs of the *P. brassicae* egg-induced response in *A. thaliana*. Treatment with PCs resulted in SA, H_2_O_2_, induction of defence genes and trypan blue staining (Stahl et al., 2020). PCs are components of cell membranes. Earlier, phospholipids of *Sogatella furcifera* were also found to induce an ovicidal response in rice (Yang, Nakayama, Toda, Tebayashi, & Kim, 2014). Our results suggest that the EAMPs in *P. brassicae* eggs that induce HR-like necrosis in *B. nigra* are other compounds than PCs. First, the *B. nigra* response is specific to *Pieris* eggs and egg wash and is absent in response to other lepidopteran eggs. Second, given our method used of washing eggs in a water-like buffer, lipids are not expected to be present in (high amounts in) the wash. Finally, as PCs are present in membranes, ovarian eggs should also induce the response. Indeed, in *A. thaliana*, ovarian eggs alone also induced *PR1* expression (Little et al., 2007). We showed that ovarian eggs did not induce HR-like necrosis.

Plant responses to insect eggs are similar to responses to (microbial) pathogens, and include SA and ROS accumulation, callose deposition, defence gene expression and cell death (Reymond, 2013). Natural variation for strength of egg-induced necrosis was found in several brassicaceous species, including *B. nigra* (Griese et al., 2021; Groux et al., 2020; Pashalidou et al., 2015). Here, we show that plants that do not express a strong HR-like necrosis, still responded with ROS accumulation and cell death as showed by trypan blue staining. Similarly, in *S. dulcamara*, variation exists for egg-induced chlorosis, and a genotype that did not respond with chlorosis and on which egg hatching rate of *S. exigua* was not reduced, still accumulated SA after oviposition (Geuss, Stelzer, Lortzing, & Steppuhn, 2017). We thus hypothesize that *Pieris* eggs generally induce an immune response in all plants of *B. rapa* and *B. nigra*, that is only in some plants accompanied by a stronger cell death response. In pathogen-induced HR, cell death can often be uncoupled from (preceding) biochemical and molecular changes and the two processes can be genetically dissected (Künstler, Bacsó, Gullner, Hafez, & Király, 2016). In that case, cell death is dispensable for resistance. However, in egg-induced HR, previous studies show that the stronger the HR-like necrosis, the higher egg mortality (Fatouros et al., 2014; Griese et al., 2021; Griese et al., 2017; Griese et al., 2020).

In *A. thaliana, PR1* expression was also found in response to crushed egg extracts of different insects (Bruessow et al., 2010; Stahl et al., 2020). In *B. nigra*, there was no cell death underneath *M. brassicae* eggs, and neither induction of *PR1* nor ethylene production in response to *M. brassicae* egg wash. Our results suggest that cell death, ethylene production and gene expression, at least in *B. nigra*, are specific to *P. brassicae* eggs, and we expect the response to be activated after detection of a Pierinae-specific elicitor. It is possible that a cellular mild defence-like response is activated against a general insect-derived EAMP, for example PCs, while the strong HR-like is activated only by EAMPs from *Pieris spp*. in those plants that can detect these. The production of ethylene after incubation with egg wash only in plants that show a strong HR-like necrosis (and not in non-HR plants), points to this effect. Mapping efforts in *B. nigra* plants, can reveal whether the genetic variation in plants is for the HR-like necrosis and/or for the detection of a *Pieris*-specific elicitor.

Molecular patterns that are detected by plants are thought to be structurally conserved molecules (Van der Burgh & Joosten, 2019). The presence of EAMPs in the glue suggests that their function could be a structural component of the glue, or a compound with an essential function to the fertilized eggs. Many proteins are found in glue of insect eggs (Li, Huson, & Graham, 2008), and egg glue of *P. brassicae* was described to consist of proteins and unsaturated lipoids (Beament & Lal, 1957). In addition, the ARG secretions could contain molecules that are produced by the parents and/or microbial symbionts to protect the vulnerable egg, for example compounds with antimicrobial activity (Flórez et al., 2018). Further identification of EAMPs can lead to new research in this direction.

In summary, we present a method to obtain EAMPs from insect eggs, and using this method, show that at least one EAMP is in the egg glue, derived from the female ARG. We furthermore assess the specificity of these elicitors and the molecular response of *Brassica* plants to *Pieris* and other eggs. The obtained knowledge paves the way for future studies on identification of EAMPs in *Pieris* egg glue, and the corresponding receptor genes in *Brassica* plants.

## Acknowledgements

We are grateful to the employees of Unifarm (WUR) for rearing and caring of the plants used in the experiment. We thank Pieter Rouweler, André Gidding, and Frans van Aggelen for rearing of *Pieris brassicae* and *Mamestra brassicae*. We thank Martijn Flipsen, Gabriella Bukovinszkine’Kiss, Klaas Bouwmeester and Arjen van der Peppel for assistance with several experiments. Guusje Bonnema and Erik Poelman are acknowledged for seeds of *B. rapa* and *B. nigra*. The authors declare having no conflict of interest.

