## Supplemental Data for "Deciphering *Brassica* plant defence responses to cabbage white butterfly egg-associated molecular patterns"

### Supplementary information file

#### Material and Methods

##### qRT-PCR

Leaf discs were snap frozen in liquid nitrogen, and RNA isolation and DNAse treatment was carried out according to the protocol by Oñate-Sánchez and Vicente-Carbajosa (2008) for vegetative tissues. For cDNA synthesis, 1µg of RNA was reverse-transcribed into cDNA using Bioline's SensiFAST cDNA synthesis kit (BIO-65054) in a 20 µl reaction volume according to the manufacturer's instructions and subsequently diluted 8 times in nuclease free water. Real time qPCR reactions were performed using Bioline's SensiFAST SYBR No-ROX Kit (BIO-98050) in 10 µl reaction volumes, containing 3 µl cDNA and 500 nM of each gene-specific primer (Supplementary Table S1) on a CFX96 Touch Real-Time PCR Detection System (Bio-Rad). The following PCR program was used for all PCR reactions: 95°C for 2 min followed by 40 cycles of 95°C for 5 s; annealing temperature for 5 s and 72°C for 10 s, with data collection at 72°C, followed by a melt curve analysis. ΔΔCq values were calculated using the Cq values of the plants prior to treatment (0 h) and normalized using the Cq values of the reference gene GAPDH.

**Supplementary Figures**

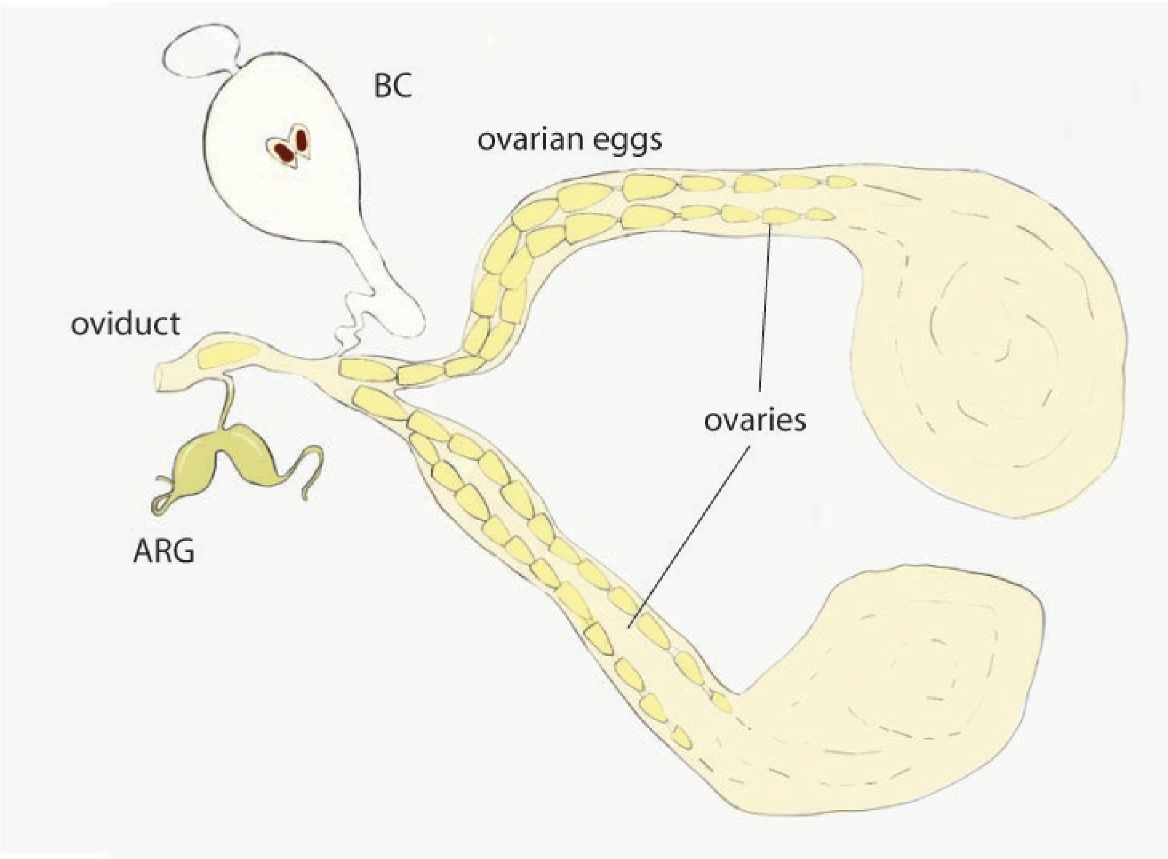

**Supplementary Figure S1**. Schematic drawing of female reproductive tract of *P. brassicae*. Pictured are the two ovary tracts, containing eggs that are developing, the bursa copulatrix (BC), the accessory reproductive gland (ARG) and oviduct. Eggs of *P. brassicae* are produced in the ovaries and then fertilized by sperm residing in the bursa copulatrix. Before eggs are oviposited onto the leaf they are covered by secretions released from the accessory reproductive gland (ARG) located right before of the oviduct.

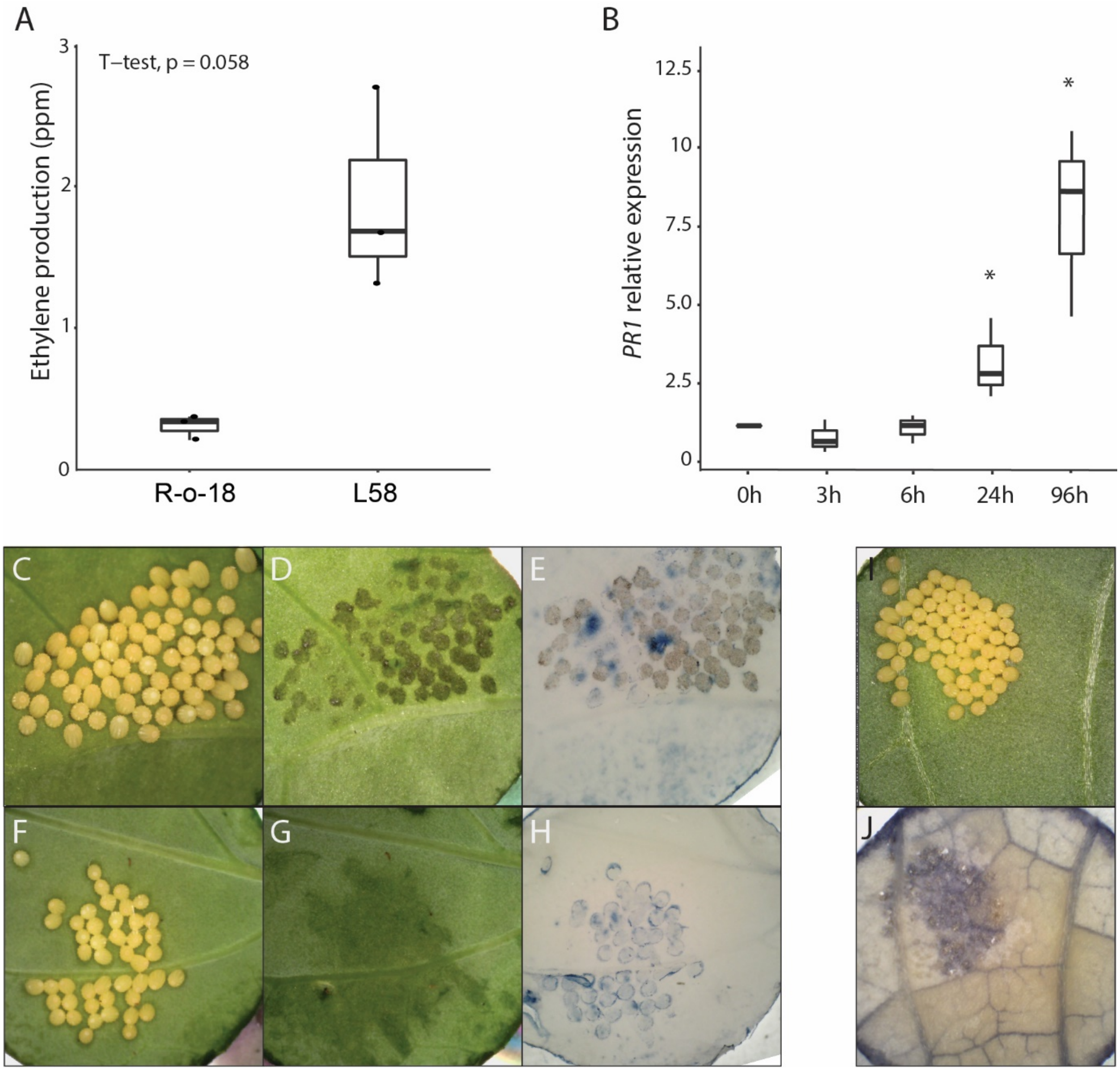

**Supplementary Figure 2**. Egg wash and eggs of *P. brassicae* induce molecular and cellular responses in *B. rapa*. A) Ethylene production in parts per million (ppm) in plants treated with egg wash in two *B. rapa* accessions. B) *PR1* expression in plants after egg deposition. Asterisks indicate significantly higher expression at the timepoint compared to the 0h timepoint (ANOVA followed by Dunnett's test, $P < 0.01$). C) *B. rapa* leaf 72 h after oviposition. D) Leaf as in (C), showing black spots underneath eggs. E) Trypan blue staining of leaf shown in (C) showing cell death underneath eggs. F). *B. rapa* leaf with no visual HR-like necrosis underneath eggs 72 hours after oviposition. G) Same leaf with eggs removed, no black spots or necrosis visible. H) trypan blue staining of leaf shown in (F) revealing dead cells underneath eggs. I) *B. rapa* leaf 24 hours after oviposition. J) NBT staining of leaf shown in (I) revealing $O_2^{\bullet -}$ deposition underneath eggs.

#### Supplementary Tables

**Supplementary Table S1:** Primer sequences and annealing temperature for primers as used in real time qRT-PCR.

| **Gene** | **Forward (5' to 3')** | **Reverse (5' to 3')** |
|---|---|---|
| ***PR1*** | CGCCGACGGACTAAGAGGCG | ACACCTCGCTTTGCCACATCCA |
| ***GADPH*** | GGAGCTGCCAAGGCTGTCGG | CCTTCAGATTCCTCCTTGATAGCC |

**Supplementary Table S2**: HR-like necrosis (score ranging from 0 to 3) expressed by *B. nigra* plants elicited by different treatments. Plants in which the eggs or egg wash tested did induce a HR-like necrosis (“HR”) and plants in which they did not (“No-HR”) are counted. Data of four separate experiments is taken together of which the mean HR frequency and HR severity is shown. Different letters indicate there was no significant difference in HR severity (Kruskal-Wallis test).

| Treatment | HR severity (SE) | HR | No-HR | HR frequency (SE) |
|---|---|---|---|---|
| **Eggs** | 2.04 (0.15) a | 33 | 18 | 0.69 (0.04) |
| **Egg wash** | 2.72 (0.17) a | 38 | 14 | 0.73 (0.04) |

**Supplementary Table S3:** Relative expression of *PR1* gene in *B. nigra* after egg deposition or treatment with egg wash. N indicates biological replicates (separate plants). HR severity is mean of all tested plants scored at 72 hours after treatment or oviposition. *PR1* relative expression is represented by mean $\pm$ se. An asterisk indicates a significant difference in mean transcript levels compared to the 0 h timepoint for eggs (ANOVA: $F_{4,14} = 6.12$, $P = 0.005$) and after treatment with egg wash (ANOVA: $F_{4,14} = 4.03$, $P = 0.022$). There were no significant differences found in mean transcript levels between the treatments for each timepoint (Student's t-tests: $P > 0.05$).

| timepoint | treatment | N | HR severity (at 72 h) | *PR1* relative expression |
|---|---|---|---|---|
| ***0 h*** | NA | 4 | NA | 1.63 $\pm$ 0.54 |
| ***3 h*** | Eggs | 4 | 2.50 | 0.67 $\pm$ 0.15 |
| | Egg wash | 4 | 3.25 | 1.47 $\pm$ 0.42 |
| ***6 h*** | Eggs | 4 | 3.00 | 55.62 $\pm$ 20.99 |
| | Egg wash | 4 | 2.75 | 5.45 $\pm$ 0.88 |
| ***24 h*** | Eggs | 4 | 3.25 | 140.34 $\pm$ 54.57 * |
| | Egg wash | 4 | 2.50 | 42.16 $\pm$ 10.54 * |
| **48 h** | Eggs | 4 | 2.75 | 131.93 $\pm$ 32.98 * |
| | Egg wash | 4 | 3.00 | 157.33 $\pm$ 39.33 |

**Supplementary Table S4:** Ethylene production after egg wash treatment in no-HR and HR plants. *Brassica nigra* leaves responded with ethylene production after incubation with egg wash for 5 hours compared to incubation with control MES buffer (Wilcoxon rank sum test, $P < 0.001$). Control treatment was an incubation of leaves in MES buffer in which the egg wash was prepared. Plant HR was determined in plants after leaves were samples for the ethylene production assay by treating another leaf with egg wash and scoring spots 72 hours after treatment. N = number of plants used for assay.

| **Plant HR** | **Mean HR severity (at 72 h)** | **treatment** | **N** | **Ethylene production (ppm)** |
|---|---|---|---|---|
| **No-HR** | 0.60 | Control | 10 | 0.000 $\pm$ 0.00 |
| | | Egg wash | 10 | 0.115 $\pm$ 0.01 |
| **HR** | 2.30 | Control | 10 | 0.000 $\pm$ 0.00 |
| | | Egg wash | 10 | 0.423 $\pm$ 0.01 |

**Supplementary Table S5**: HR- like necrosis (score ranging from 0 to 3) expressed by *B. nigra* plants elicited by different treatments. . Plants in which egg wash tested did induce a HR-like necrosis ("HR") and plants in which they did not ("No-HR") are counted. Different letters indicate differences in HR frequency (GLM) or HR severity (Kruskal-Wallis test).

| Treatment | HR severity (SE) | HR | No-HR | HR frequency (SE) |
|---|---|---|---|---|
| **Egg wash – eggs 1 day old** | 2.5 | 20 | 6 | 0.77 |
| **Egg wash – eggs 2 days old** | 2.38 | 10 | 3 | 0.77 |
| **Egg wash – eggs 3 days old** | 2.31 | 9 | 4 | 0.69 |
| **Egg wash – eggs 4 days old** | 2.15 | 8 | 5 | 0.62 |
| **Egg wash – eggs 5 days old** | 2.08 | 9 | 4 | 0.69 |
| **Hatched eggs** | 1.84 | 8 | 5 | 0.62 |
| **Egg wash** | 3.38 (0.20) | 6 | 2 | 0.75 |
| **Unfertilized egg wash** | 2.75 (0.04) | 20 | 4 | 0.83 |
| **Egg wash** | 3.6 (0.18) | 5 | 0 | 1.00 |
| **ARG** | 3.6 (0.11) a | 5 | 0 | 1.00 |
| **Ovarian eggs** | 0.2 (0.08) b | 0 | 5 | 0.00 |
| **Bursa copulatrix** | 0.8 (0.08) b | 0 | 5 | 0.00 |
| **ARG unmated** | 3.8 (0.08) a | 5 | 0 | 1.00 |
| **Egg wash** | 3.0 (0.05) a | 15 | 1 | 0.94 |
| **Wash glue** | 2.5 (0.08) a | 10 | 2 | 0.83 |
| **Glue removed (bleach)** | 1.0 (0.00) b | 0 | 8 | 0.00 |
| **Glue removed (NaPO4)** | 1.0 (0.07) b | 1 | 7 | 0.13 |

**Supplementary Table S6**: Relative expression of *PR1* gene in *B. nigra* 24 hours after treatment with either *P. brassicae* or *M. brassicae* egg wash. Control is a treatment with MES buffer in which the egg wash was prepared. N indicates biological replicates (separate plants). *PR1* relative expression is represented by mean ± se. Different letters indicate significant differences in mean *PR1* expression, ANOVA followed by Tukey, P<0.001.

| **Egg wash** | **N** | ***PR1*** **relative expression** |
|---|---|---|
| ***Control*** | 5 | 1.24 ± 0.19 |
| ***Pieris brassicae*** | 4 | 159.48 ± 14.39 |
| ***Mamestra brassicae*** | 4 | 4.18 ± 0.80 |
